# Direct evidence of an increasing mutational load in humans

**DOI:** 10.1101/307058

**Authors:** Stéphane Aris-Brosou

## Abstract

The extent to which selection has shaped present-day human populations has attracted intense scrutiny, and examples of local adaptations abound. However, the evolutionary trajectory of alleles that, today, are deleterious has received much less attention. To address this question, the genomes of 2,062 individuals, including 1,179 ancient humans, were reanalyzed to assess how frequencies of risk alleles and their homozygosity changed through space and time in Europe over the past 45,000 years. While the overall deleterious homozygosity has consistently decreased, risk alleles have steadily increased in frequency over that period of time. Those that increased most are associated with diseases such as asthma, Crohn disease, diabetes and obesity, which are highly prevalent in present-day populations. These findings may not run against the existence of local adaptations, but highlight the limitations imposed by drift and population dynamics on the strength of selection in purging deleterious mutations from human populations.

## Introduction

The role played by natural selection in shaping present-day human populations has received extensive scrutiny (1; 2; 3), especially in the context of local adaptations (4). However, most studies to date assume, either explicitly or not, that populations have been in their current locations long enough to adapt to local conditions (5), and that their effective population sizes were large enough to allow for the action of selection (6). If these conditions were satisfied, not only should selection be effective at promoting local adaptations, but deleterious alleles should also be eliminated over time.

Surprisingly, this question has only been examined by comparing present-day modern humans from Africa and Europe (7), leading to the consensus that selection was as effective in Europeans as in Africans (8; 9) but without really assessing the effectiveness of selection on a large data set (10). Here, I examine this question directly, based on the genotype of not only present-day humans, but also of > 1, 000 ancient humans over the past 45,000 years.

## Results and Discussion

In order to trace the evolutionary trajectory of deleterious alleles in human populations over time, the genotypes of 2,062 individuals, of which 1,179 were ancient humans, were assessed at about 1.2 million Single Nucleotide Polymorphisms (SNPs; the ‘1240k capture’ (11)). The present study focuses on individuals sampled throughout Europe, from the Atlantic to the Ural Mountains, as this is where most ancient DNA has been recovered and genotyped throughout the world. To take into account the history of admixture and replacement events characterizing Europe (12; 11), this region was split into four quadrants along two orthogonal axes roughly cutting through the Alps (Figure 1). These four quadrants represent approximately four distinct areas that played significant roles in the three peopling waves that affected Europe, first from the Fertile Crescent (Q4 in Figure 1) to the rest of Europe during the Paleo-/Mesolithic period, between 45 and 10 thousands years ago (kya) (13; 14). These early Europeans were then replaced after the Last Glacial Maximum, 10 kya, by Neolithic farmers (15) who reached the Iberian Peninsula (Q3), and Britain and Scandinavia (Q2) 7-6 kya (16). A third major migration event swept into Europe, including central Europe (Q1), during the late Neolithic and the early Bronze Age, about 3.5 kyr ago (11; 17). This splitting scheme is a simplification that has the advantage of relying solely on the geotagging of each individual, while approximately reflecting their shared ancestry. Under the constraint of analyzing approximately even numbers of individuals across these four quadrants, the timing of these migration events suggested splitting Europe into four different time periods, representing roughly the Mesolithic (t1: 45-10 kya), the Neolithic (t2: 10-3.5 kya), the Bronze Age (t3: 3.5-0 kya), and present-day populations (t4: 0-present) as in Figure 1.

**Figure 1.**
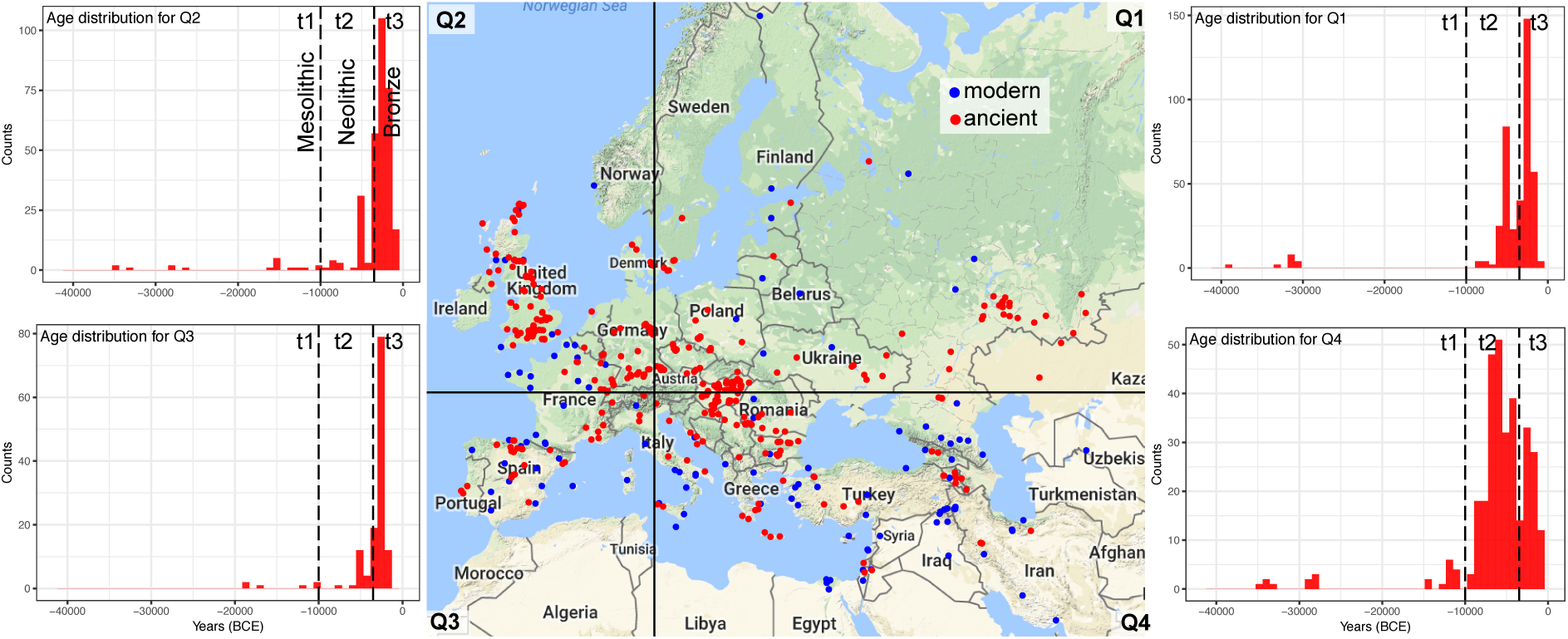
Spatiotemporal distribution of the human genomes analyzed in this study. Center: Sampling location of modern (blue) and ancient (red) individuals throughout Europe. The maps is divided into four quadrants representing northeastern (Q1), northwestern (Q2), southwestern (Q3), and southeastern (Q4) Europe. By each quadrant, the age distributions of the ancient humans are shown; the four time periods considered correspond roughly to the mesolithic (t1: 45,000–10,000 BCE), the neolithic (t2: 10,000–3,500 BCE), the bronze age (t3: 3,500–0 BCE), and modern humans (t4: the 2000 CE). Map background: Google (2018).

To find which genomic positions among these 1.2 million SNPs are deleterious, a database of markers identified based on genome-wide association studies (DisGeNET) was used (18). This database contains 83,582 SNPs, and the associated risk alleles were retrieved separately. Because these SNPs were discovered in present-day populations, the results that follow are conditional on the fact that the derived mutations at these SNPs survived until now, and have not yet fixed. Strikingly, the average frequency of deleterious mutations significantly increased through each time period in all four quadrants (Figure 2). One possible explanation for this nonadaptive pattern is that most of the SNPs tested are weakly deleterious, and increased in frequency at the same rate as nearly-neutral mutations would have under drift (19).

**Figure 2.**
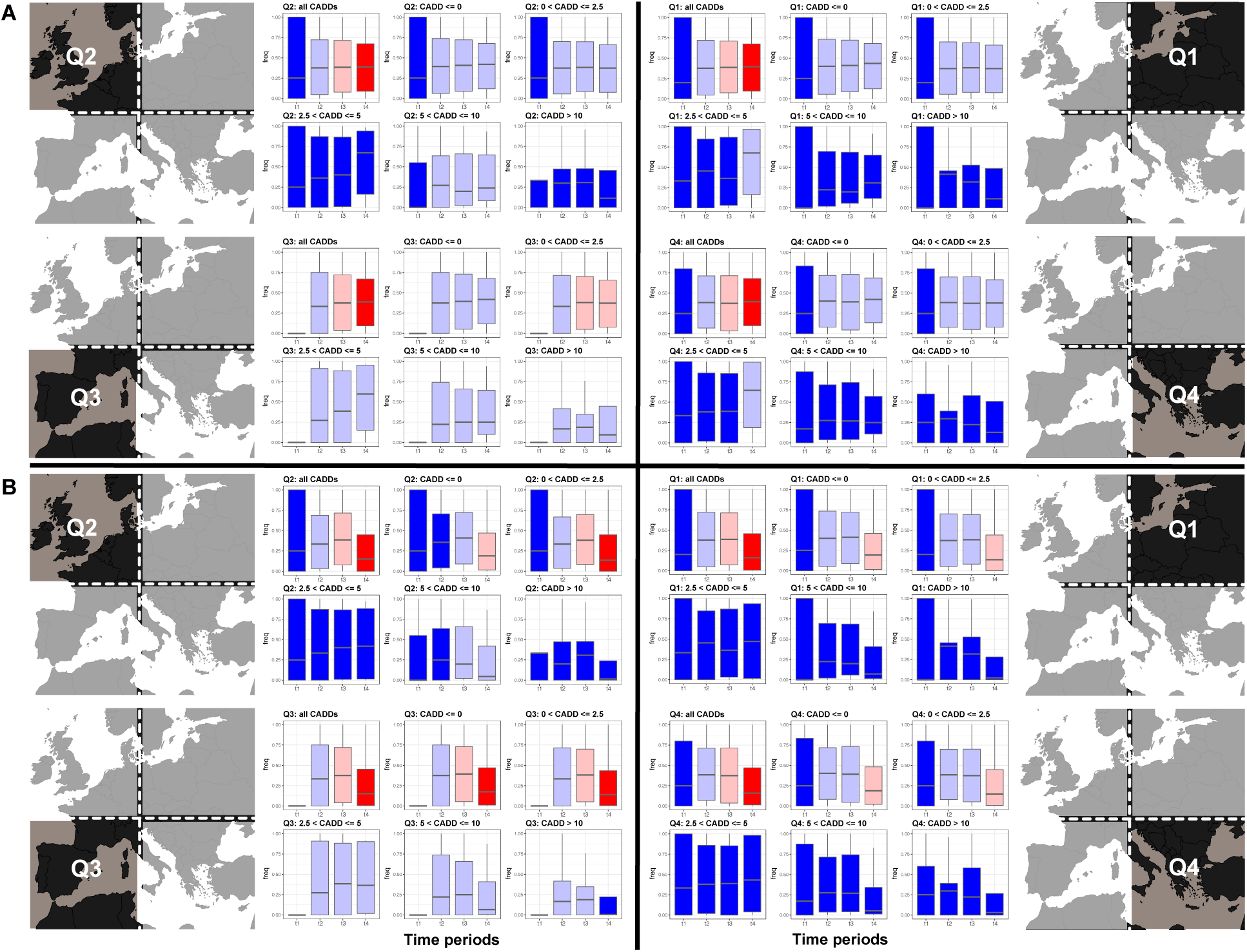
Evolution of SNP frequencies through space and time. (**A**), Frequencies of risk alleles. (**B**), Homozygosity at risk alleles. The top left panel in each quadrant show the frequency distributions at all SNPs over the four time periods, while the next five panels show the frequency distributions at SNPs in each severity category, ranging from benign (CADD < 0) to severe (CADD ≥ 10) effects. Color differences in each panel indicate significance of the Dunn test at the 1% level with Benjamini-Hochberg correction for multiple tests – colors are ordered from significantly lower (cooler) to significantly higher (warmer) frequencies of the risk alleles.

To test if weakly deleterious mutations that survived until now are driving the increase in risk allele frequency, their relative pathogenicity was assessed by the Combined Annotation Dependent Depletion (CADD) score (20), which take values from −6 (the least deleterious) to 100. The values observed here range from about −5 to 15, are sharply distributed around 0, and are skewed to the right (Figure S1). To establish a baseline among the observed CADD scores, their range was split into five bins. Among those, the mutations that are the most weakly deleterious (up to CADD = 0) are also the most likely to be nearly neutral, and can thus be used as a control. In all four quadrants, these most weakly deleterious mutations significantly increased in frequency from t1 to t2 (Figure 2A). As these mutations are the most numerous (Figure S1), they are most likely responsible for the initial increase in frequency, between t1 and t2. However, at higher CADD scores, significant increases are observed after the t1/t2 transition, in particular between t2 and t3 (Q3), and t3 and t4 (Q1, Q4). Along with the overall pattern of continuous increase from t1 to t4 (Figure 2A), these results show that some deleterious mutations increased in frequency, and were able to do so beyond the variation observed at our control mutations, the most weakly deleterious. Now if the mutations with a high CADD score were recessive, these deleterious alleles would mostly be present in the heterozygous genotypes, and their homozygosity would be low.

To indirectly assess the effect of dominance on the general pattern of increasing frequency of risk alleles through time, the homozygosity of the risk alleles was calculated (Figure 2B). Overall, homozygosity decreased significantly through time, but again, this pattern is mostly driven by the least deleterious alleles (up to CADD = 0). The most deleterious ones (CADD > 10), on the other hand, saw their heterozygosity decrease recently (in t4) only in southwestern Europe (Q3). In all the three other quadrants, the heterozygosity of the most deleterious alleles did not change significantly (Figure 2B), suggesting that these alleles are only partially recessive.

To better understand which diseases increase in frequency through time, robust linear models were fitted to each SNP, irrespective of its CADD score. Because GWAS are most powerful at identifying associations based on common variants (21), the SNPs deposited in DisGeNET, and used here, are at intermediate frequencies in present-day populations (Figure S2A), and the SNPs with extreme allele frequencies have indeed a smaller chance to show significant change in frequency through time (Figure S2B). From the volcano plot corresponding to these regressions of risk allele frequency again time (Figure 3A), two groups of SNPs can be defined: those whose frequency significantly decreases (top left area of volcano plot) or increases (top right area) through time. Both groups show an increase in slope with present-day allele frequency (Figure S3), mirroring the two sides of the volcano plot. Assuming that older alleles have higher frequencies (22), this result suggests that: (i) for risk alleles that decrease in frequency, older alleles disappear slower than more recent variants; (ii) for risk alleles that increase in frequency, older alleles increase in frequency faster than more recent ones. This pattern may be due the GWAS bias highlighted above (Figure S2), but could also reflect the differential strength of selection through time, being less effective in the past than in more recent times. Tentative support for this hypothesis comes from the variation of CADD scores with the age of risk alleles (frequency in present-day population): although none of this patterns are significant, the trends suggest that (i) for risk alleles that decrease in frequency, older alleles have higher CADD scores (are more deleterious; Figure S4A), while (ii) for risk alleles that increase in frequency, older alleles have lower CADD scores (Figure S4B). More detailed reconstructions might be warranted to further test this hypothesis.

**Figure 3.**
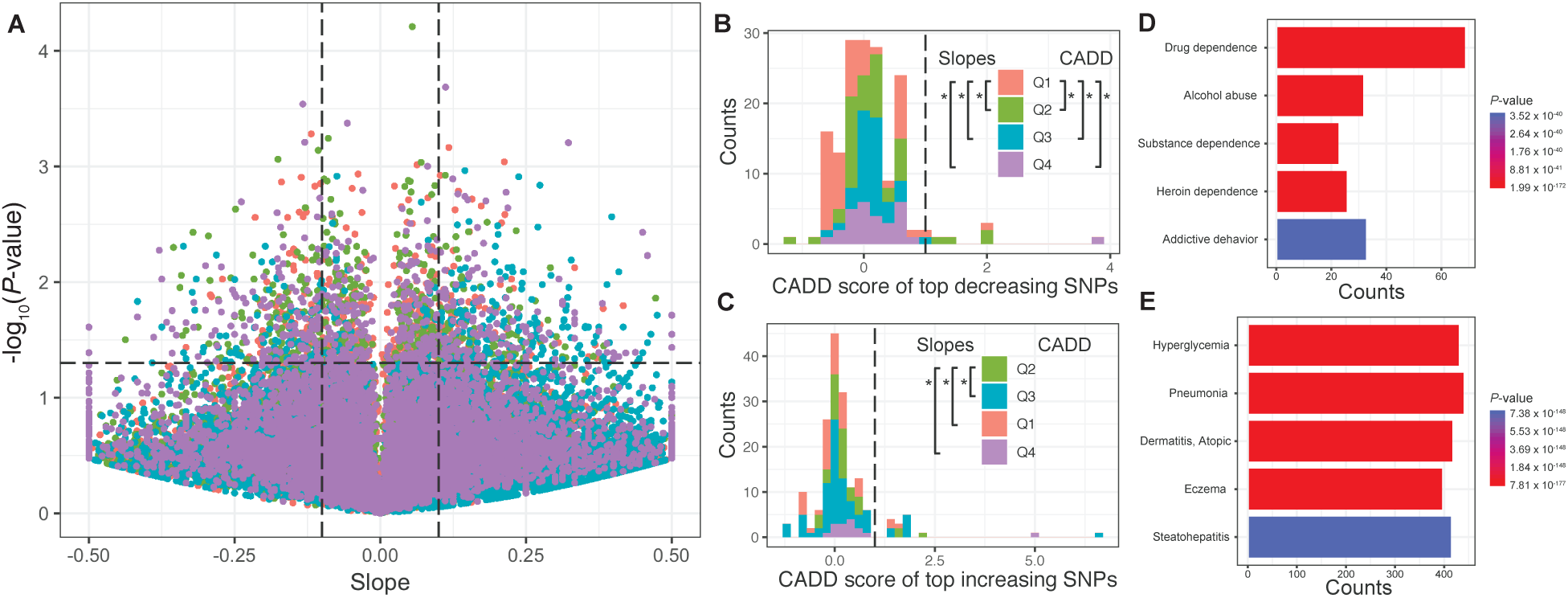
Volcano plot of the regressions of SNP frequencies through time. (**A**), The plot shows the *P*-values, transformed on a − log_10_ scale, as a function of the slope estimates of the robust regressions, with the four quadrants color coded (Q1: salmon, Q2: green, Q3: blue, Q4: magenta). (**B**), Distribution of the CADD scores for the SNPs whose frequency decreases through time (*P* ≤ 0.05, slope ≥0.1). The Dunn test was used to assess differences between slopes (left of inset legend) and CADD scores (right of inset legend) at an FDR ≤ 0.05. (**C**), Same as in (**B**) for the SNPs whose frequency increases through time. Top five enriched gene names among the top decreasing (**D**), and top increasing (**E**) genes identified from the volcano plot with CADD scores ≥ 1.

Irrespective of temporal changes in the strength of selection, in the first group (risk alleles that decrease in frequency), quadrant Q1 has both the most negative slopes and the smallest CADD scores (Figure 3B; Dunn test, FDR < 0.05), and the most affected phenotypes, with a CADD score greater than 1, include “age at menarche” and “drug dependence” (Table S1). In the second group (risk alleles that increase in frequency), quadrant Q2 has the most positive slopes even if all quadrants has similar CADD scores (Figure 3C; Dunn test, FDR < 0.05), and the most affected phenotypes, with a CADD score greater than 1, include “asthma,” “Crohn disease,” “Diabetes Mellitus” and “obesity” (Table S1) – diseases that tend to be frequent in present-day populations of European descent. Enrichment analyses confirmed that SNPs involved in addictive behavior are overrepresented in the set of SNPs with decreasing frequency, while common diseases are overrepresented in the group with increasing frequencies. Note that in both sets of SNPs, the CADD scores are never greater that 10, suggesting that the most deleterious alleles either do not change in frequency, and / or do so at a very slow rate – which again is consistent with their not being eliminated through time.

The efficacy of selection at removing deleterious mutations depends on fitness effects, with the most deleterious variants being eliminated with the greatest ease. However, this efficacy can be affected by a number of factors such as the effective population size, which is arguably the most critical (23). Indeed, ancient human demographics is marked during the Last Glacial Maximum (LGM) by a severe bottleneck, which left traces in present-day individuals of European ancestry (24) as well as in other worldwide populations (25). The results presented here are therefore consistent with the population genetics and demographic contexts, which permitted the nonadaptive increase in frequency over time of deleterious variants.

Although the variance of deleterious allele frequencies in the oldest time period (t1: the Mesolithic) is very large, their median is also quite high (∼ 0.25), begging the question of the origin of these variants. Previous work showed that some of these deleterious variants found in modern humans could have a Neanderthal origin, and it was quickly suggested that selection could be efficient at removing these variants (26), especially in East Asians, as selection is mostly acting on weakly deleterious mutations located in protein-coding genes (27). However, recent work showed that the decline of Neanderthal ancestry in modern humans was likely due to demographic events such as migrations between West Eurasian and African populations, rather than widespread and systematic selection against these alleles (28). This is entirely consistent with the results presented here, where frequencies of even the most weakly deleterious variants do not significantly change over time. This is potentially due to most of the SNPs used here being located out-side of protein-coding genes – which again is consistent with the depletion of Neanderthal alleles in regulatory and conserved noncoding region (28).

An allelic frequency however represents a snapshot of a dynamic process, where variants are generated by mutations, and eliminated by selection (or lost by drift). The initial increase, between the Mesolithic and the Neolithic (t1 and t2, respectively), could be due to drift, but also to an increased mutational pressure. Little evidence however seems to support this latter possibility, except maybe for recent times, where humans tend to expose themselves to mutational agents, leading to a relaxation of selection against mildly deleterious mutations (29). As a result, nonadaptive factors following the initial LGM bottleneck remain the most plausible explanation for the monotonic increase in frequency of deleterious alleles in the European human population.

Again, this is entirely consistent with the demographic context, as this geographic region has been affected by at least three migration waves, associated with rapid demographic expansion(-s) (24). Indeed, range expansions have been shown to enable new deleterious mutations to behave almost like neutral variants, hereby facilitating their increase in frequency (30). This would suggest that (i) the deleterious mutations observed post-LGM are novel, *i*.*e*. do not have a Neanderthal origin, and (ii) increase in frequency because of a nonadaptive factor (repeated range expansions). One signature that is missing from typical range expansions is a frequency gradient along the expansion axis (from Q4 to the other quadrants; 31); this absence of gradient may however be caused by multiple expansions, whose individual directions are not necessarily overlapping. This may also be the reason why an increase in the homozygosity of deleterious variants was not observed here. As a result, the existence of geographic regions of high fitness in which the effects of deleterious mutations would be partially masked, as predicted by range expansion models (32), seems unlikely in European humans.

The assessment of the phenotypic effects of variants was based on CADD scores, which correlate with a number of experimental data related to pathogenicity, disease severity, *etc*. (20), but that also show a number of discrepancies in select cases (33; 34), or in a study of 12,391 cancer-related SNPs (35). While this number represents 12% of the SNPs analyzed above, it is unlikely that the results presented here are affected by the predictive power of CADD scores. Indeed, previous studies used a different score, the Genomic Evolutionary Rate Profiling (GERP) Rejected Substitution (RS) score, to predict the phenotypic impact of mutations (36), and showed either the existence of a spatial increase of deleterious mutations from Africa to the rest of the world, consistent with the expansion axes out of Africa (25), or an increase of deleterious alleles at the front of a range expansion compared to the inner core of the source population (37). The data presented here add a temporal dimension to these previous spatial results, which were found at a broader scale. It is therefore possible that the results presented here, limited to Europe, be generalizable to the rest of the non-Subsaharan human populations. Simulations can be useful by shedding light on some of the processes affecting selection (8; 9), but are usually limited to simple scenarios (28) that are no substitute for actual data from the past. As more ancient DNA from the rest of the world is accumulating (38; 39; 40), it will soon be possible to test the effectiveness of selection at purging deleterious mutations from non-Subsaharan human populations.

## Materials and Methods

### Genotypic Data

Complete genotypic data were retrieved for nine ancient DNA studies that mostly focused on European individuals (12; 11; 41; 42; 43; 44; 45; 46; 47) from the David Reich website (reich.hms.harvard.edu). The ‘geno’ files were first converted into ‘ped’ format with EIGENSTRAT v.6.1.4 (48), and the ‘map’ files that contain information relative to SNP identifiers were created with a custom R script (see github.com/sarisbro for all the scripts developed in this study). The human reference genome, as well as those of the Neanderthal, Denisovan, and the Chimpanzee were removed. The date and sampling co-ordinates of each of the 3,409 unique individual were extracted from the Supplementary Information files of each publication (Data S1). Individuals from outside of Europe were eliminated to keep only those north of the Arabian peninsula and south of the Arctic (latitudes ∈ [20°, 80°]), and west of Iceland / Cabo Verde and east of Mount Ural (longitudes ∈ [−10°, 60°]). This European region was further divided into four quadrants Q1 to Q4, placing the crosshair at 12°W and 47°N, so that each region has a similar number of individuals in each time period t1 to t4 (present-day individuals: 126 in Q1; 67 in Q2; 166 in Q3; 524 in Q4: total present-day 883; ancient individuals: 410 in Q1; 315 in Q2; 134 in Q3; 320 in Q4: total ancient 1,179). These time periods correspond roughly to the mesolithic (t1: 45,000–10,000 BCE), the neolithic (t2: 10,000–3,500 BCE), the bronze age (t3: 3,500–0 BCE), and modern humans (t4: the 2000s CE).

Each individual was genotyped at 1,233,013 SNPs (the ‘1240k capture’, (11)), and hence contained information for twice as many or 2,466,026 alleles. To avoid computational issues related to loading large files (with 2,466,026 rows for 2,062 individuals) into memory, each genome was saved to an individual file.

### Phenotypic Data

Disease data were retrieved from DisGeNET (18) at www.disgenet.org/ds/DisGeNET/results/curated_variant_disease_associations.tsv.gz, which contained 83,582 SNP identifiers involved in genetic diseases. The genotype of each these 83,582 SNPs was then retrieved from the 2,062 individuals, binned within their quadrant and time period (Data S2). Risk alleles were retrieved from SNPedia (49) and SelfDecode (www.selfdecode.com; Data S3). Allele and genotype frequencies were calculated by counting non-missing alleles, and dividing each count by the total number of allele counted, or half that number, respectively (Data S4).

Severity of each SNP was assessed not by any of the DisGeNET scores, that are some-what *ad hoc*, but by their Combined Annotation Dependent Depletion (CADD) score (20). For this, a list of all possible SNVs in GRCh37/hg19 was downloaded from the CADD web-site (krishna.gs.washington.edu/download/CADD/v1.3/whole_genome_SNVs.tsv.gz), split into multiple files containing 100 million lines each (to avoid loading a single 250GB text file in memory), from CADD score was extracted at each SNPs in DisGeNET (Data S4).

### Matching Genotypic and Phenotypic Data

In each quadrant and each time period, a list of corresponding individual was extracted, and used to estimate the frequency of the risk allele in each quadrant / time period combination. Significant frequency differences were assessed with the Dunn test, with Benjamini-Hochberg correction for multiple comparisons at a FDR ≤ 0.05 unless otherwise stated. Robust linear regressions were used to find SNPs with decreasing and increasing frequency through time. Enrichment analyses for these decreasing and increasing SNPs were done with the clusterProfiler R package (50), based on lists of gene names containing the SNPs of interest, with respect to all gene names with SNPs in DisGeNET. All analyses were performed with R (51) ver. 3.2.3.

## Acknowledgements

I would like to thank first and foremost David Reich for making all the genotyping data produced in his lab freely available online. My gratitude also goes to Kelly Burkett for providing extensive comments, as well as to Joseph Lachance and Yassine Souilmi for discussions. This work was partly conducted while being at the University of Hokkaido in Hakodate, thanks to an Invitational Fellowship from the Japanese Society for the Promotion of Science, and was supported by the Natural Sciences Research Council of Canada. It would not have been possible without assistance from the Center for Advanced Computing and Compute Canada.

## 1 Supplementary Figures

**Figure S1.**
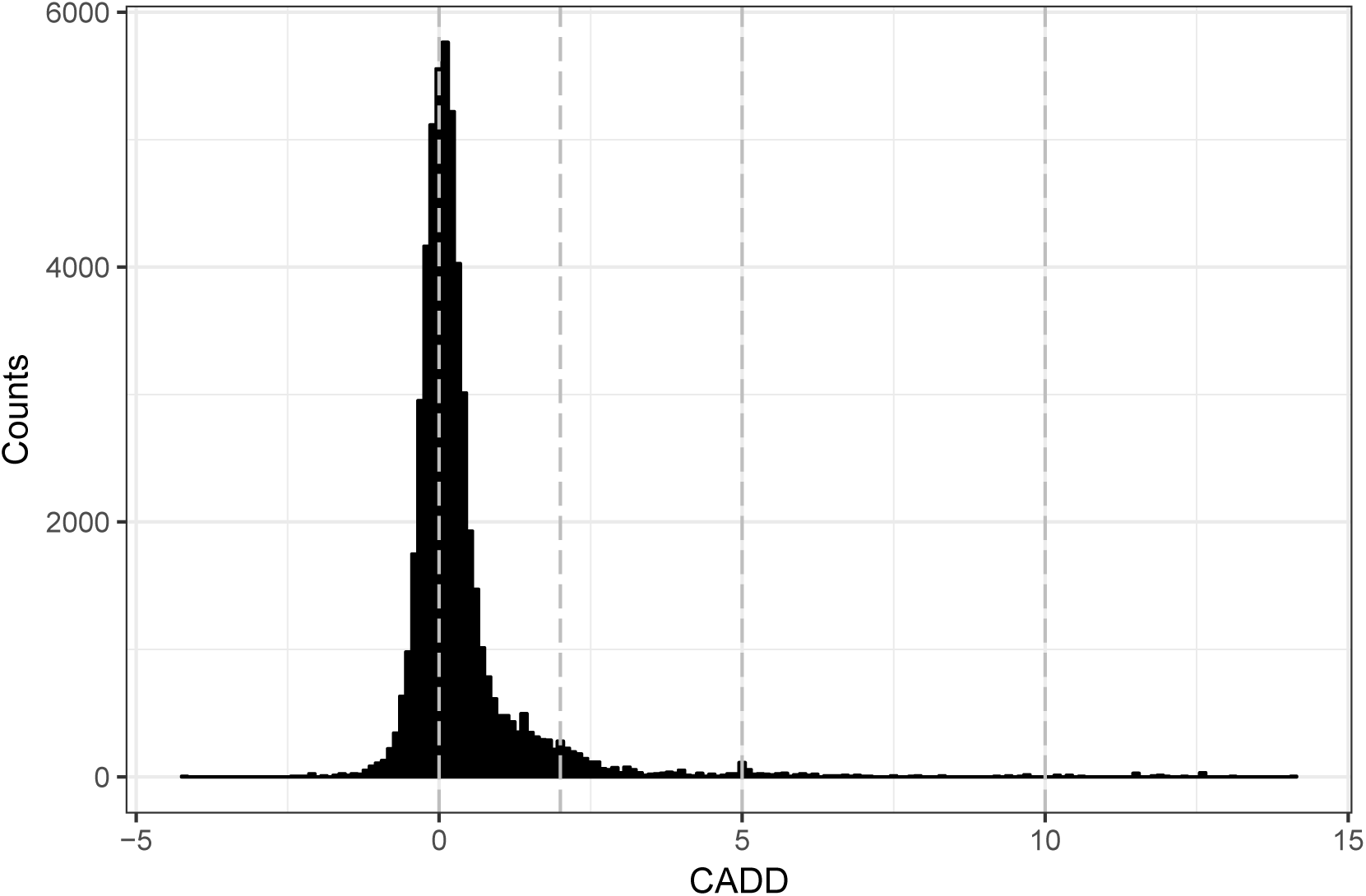
Distribution of CADD scores for the SNPs in DisGeNET. CADD scores are shown for the 83,582 SNPs contained in DisGeNET 4.0 database. Vertical lines split SNPs into five categories ranging from inferred to have benign effects (CADD < 0) to severe effect (CADD ≥ 10).

**Figure S2.**
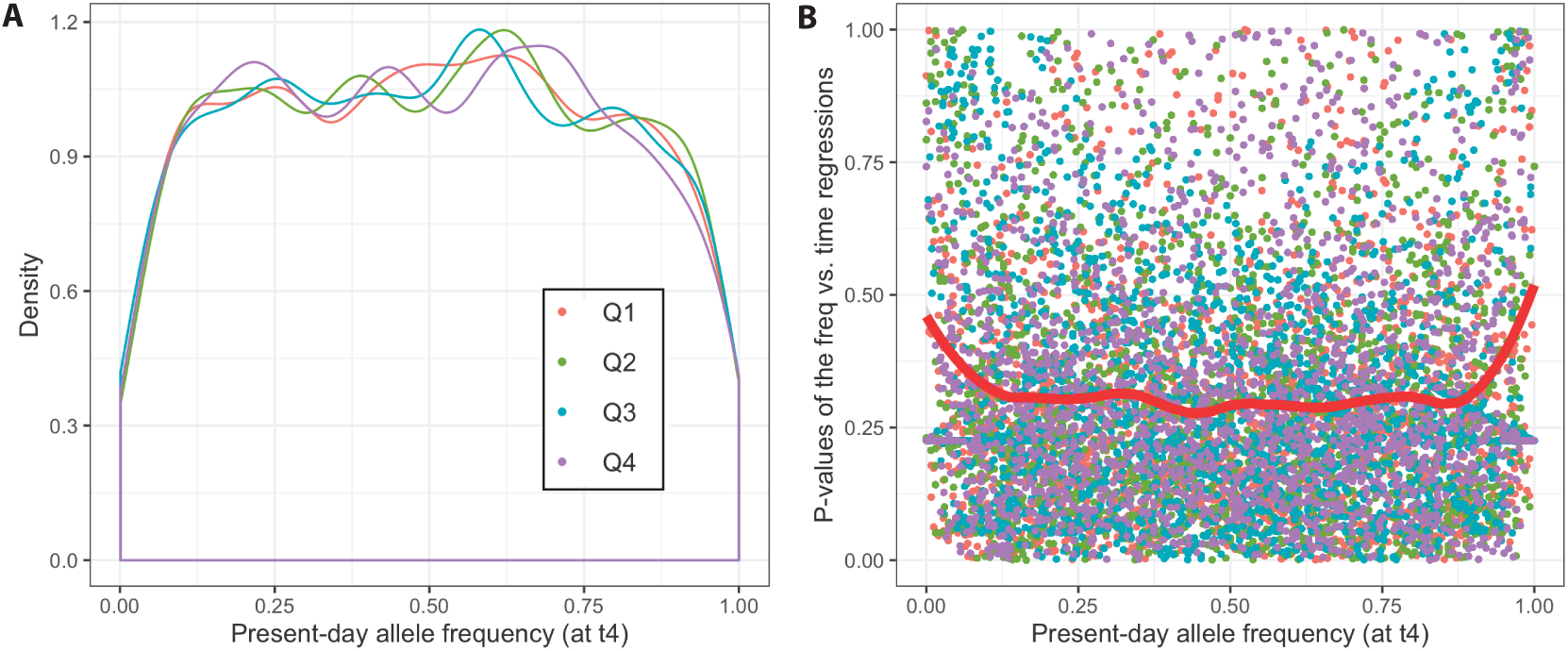
GWAS data are enriched in SNPs of intermediate frequency. (**A**), density of the frequency of the SNPs used in this study; note the paucity of SNPs at both ends of the frequency spectrum. (**B**), significance of the regressions as in Figure 3A. The fitted loess curve is shown in red. In both panels, quadrants are color-coded as shown in panel (**A**).

**Figure S3.**
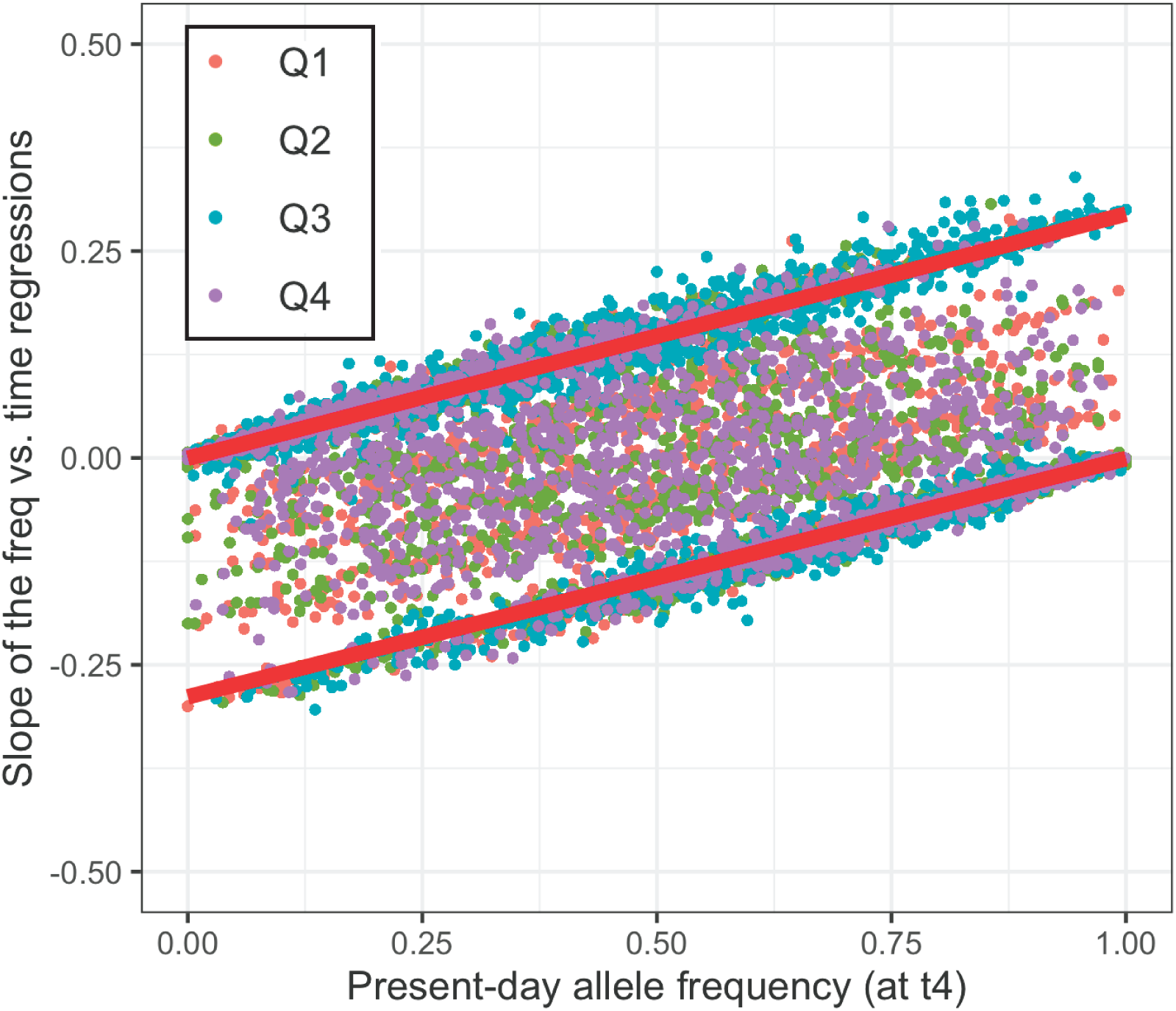
Slopes of the regressions as in Figure 3A as a function of the frequency of risk alleles at time t4 (present day). Robust regression lines (in red) are shown for negative (lower line; *t* = 3, 395, *df* = 835,497, *P* < 2 × 10^−16^) and positive (upper line; *t* = 3, 821, *df* =996,666, *P* < 2 × 10^−16^) slopes.

**Figure S4.**
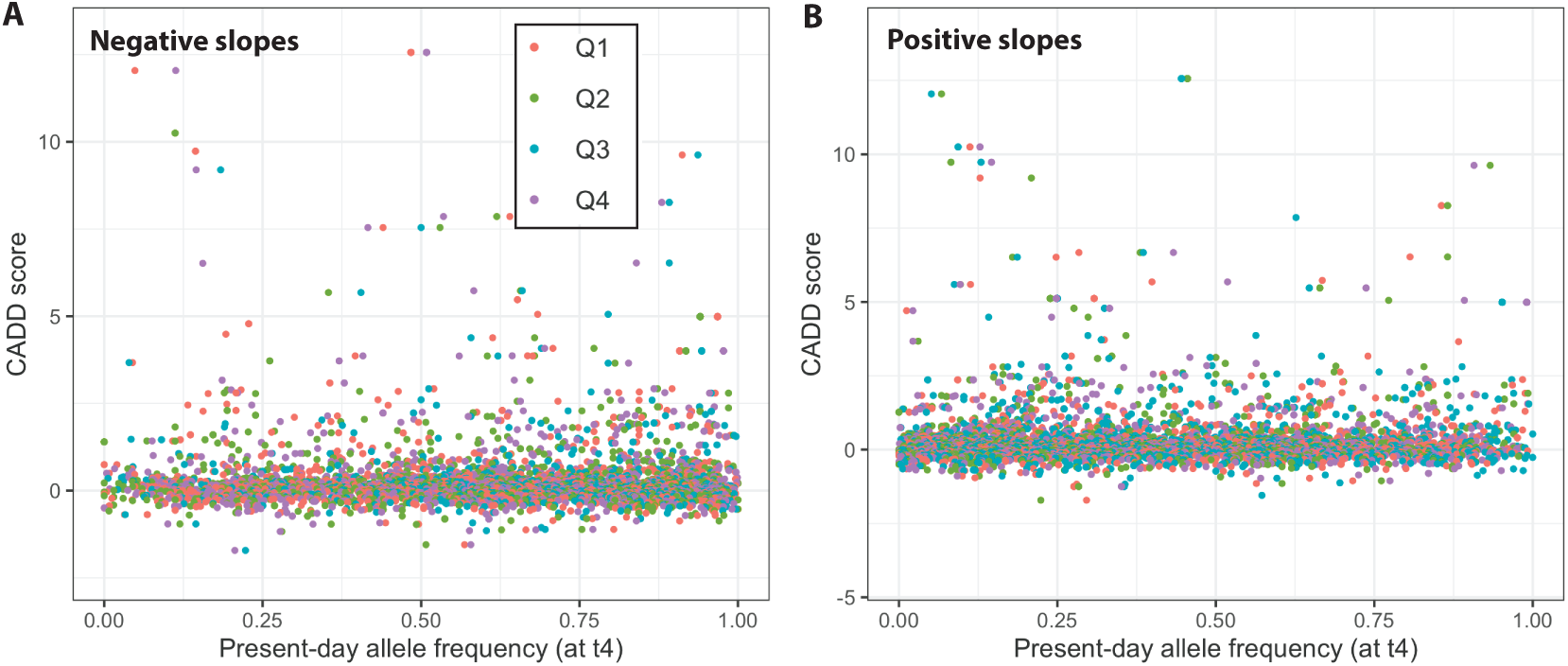
Present-day frequency of risk alleles and CADD scores. Slopes of the regressions of risk allele frequencies against time (as in Figure 3A). (**A**), for the negative slopes (*t* = 1.747, *df* = 4674, *P* = 0.0808). (**B**), for the positive slopes (*t* = −0.480, *df* = 5524, *P* = 0.6310). In both panels, quadrants are color-coded as shown in panel (**A**).

## 2 Supplementary Table

**Table S1.** Sets of SNPs and their associated phenotype that are significantly decreasing (top) and increasing (bottom) through time. For each SNP, the table shows the quadrant ID, the SNP frequency at times t1 to t4, the SNP identifier, the disease identifier and name from DisGeNET, the CADD score and the slope and *P*-value from the robust regression of SNP frequency *vs*. time.

| Decreasing | Quad ID | t1 | t2 | t3 | t4 | SNP ID | Disease ID | Disease name | CADD | Slopes | $P$-values |
| --- | --- | --- | --- | --- | --- | --- | --- | --- | --- | --- | --- |
|  | Q4 | 1.0000 | 0.7532 | 0.7059 | 0.5602 | rs1052030 | C1568247 | Usher Syndrome Type I | 3.8589 | -0.1367 | 0.0355 |
|  | Q2 | 1.0000 | 0.8462 | 0.7303 | 0.6970 | rs807601 | C0201976 | Creatinine measurement serum | 2.0105 | -0.1025 | 0.0337 |
|  | Q2 | 1.0000 | 0.8462 | 0.7303 | 0.6970 | rs807601 | C1561549 | Glomerular filtration rate finding | 2.0105 | -0.1025 | 0.0337 |
|  | Q1 | 1.0000 | 0.6582 | 0.5660 | 0.3577 | rs951366 | C1314691 | Age at menarche | 1.9794 | -0.2019 | 0.0263 |
|  | Q2 | 1.0000 | 0.8182 | 0.6923 | 0.5909 | rs12995333 | C1510472 | Drug Dependence | 1.4576 | -0.1353 | 0.0090 |
|  | Q2 | 1.0000 | 0.7750 | 0.5955 | 0.5149 | rs913588 | C1314691 | Age at menarche | 1.1493 | -0.1635 | 0.0195 |
| Increasing | Quad ID | t1 | t2 | t3 | t4 | SNP ID | Disease ID | Disease name | CADD | Slopes | $P$-values |
|  | Q3 | 0.0000 | 0.0833 | 0.2745 | 0.3855 | rs2304725 | C0004096 | Asthma | 6.6726 | 0.1348 | 0.0106 |
|  | Q4 | 0.5000 | 0.7034 | 0.7727 | 0.8922 | rs3816183 | C0848558 | Hypospadias | 5.0550 | 0.1246 | 0.0216 |
|  | Q2 | 0.0000 | 0.0882 | 0.2875 | 0.2955 | rs604723 | C0428886 | Mean blood pressure | 2.2455 | 0.1086 | 0.0487 |
|  | Q3 | 0.0000 | 0.3333 | 0.8889 | 0.9458 | rs11209026 | C0009324 | Ulcerative Colitis | 1.7046 | 0.3393 | 0.0370 |
|  | Q3 | 0.0000 | 0.3333 | 0.8889 | 0.9458 | rs11209026 | C0010346 | Crohn Disease | 1.7046 | 0.3393 | 0.0370 |
|  | Q3 | 0.0000 | 0.3333 | 0.8889 | 0.9458 | rs11209026 | C0021390 | Inflammatory Bowel Diseases | 1.7046 | 0.3393 | 0.0370 |
|  | Q3 | 0.0000 | 0.3333 | 0.8889 | 0.9458 | rs11209026 | C0033860 | Psoriasis | 1.7046 | 0.3393 | 0.0370 |
|  | Q3 | 0.0000 | 0.3333 | 0.8889 | 0.9458 | rs11209026 | C0038013 | Ankylosing spondylitis | 1.7046 | 0.3393 | 0.0370 |
|  | Q2 | 0.2000 | 0.4318 | 0.6000 | 0.6642 | rs507506 | C2700366 | Adiponectin Measurement | 1.5824 | 0.1561 | 0.0280 |
|  | Q1 | 0.0000 | 0.0513 | 0.2018 | 0.3306 | rs12521868 | C0010346 | Crohn Disease | 1.5272 | 0.1142 | 0.0167 |
|  | Q2 | 0.0000 | 0.0625 | 0.2080 | 0.3769 | rs12521868 | C0010346 | Crohn Disease | 1.5272 | 0.1276 | 0.0180 |
|  | Q1 | 0.0000 | 0.1800 | 0.2568 | 0.3387 | rs2617170 | C0004943 | Behcet Syndrome | 1.4717 | 0.1093 | 0.0241 |
|  | Q3 | 0.0000 | 0.2500 | 0.3571 | 0.5934 | rs1421085 | C0011860 | Diabetes Mellitus Non-Insulin-Dept | 1.3791 | 0.1887 | 0.0104 |
|  | Q3 | 0.0000 | 0.2500 | 0.3571 | 0.5934 | rs1421085 | C0028754 | Obesity | 1.3791 | 0.1887 | 0.0104 |
|  | Q1 | 0.3333 | 0.5000 | 0.6173 | 0.6613 | rs9854771 | C0235974 | Pancreatic carcinoma | 1.3591 | 0.1101 | 0.0299 |

## 3 Supplementary Data

Supplementary data are available at:

https://github.com/sarisbro/data/tree/master/AncientHumans

**Data S1. Individuals used in this study**. List of individuals used in this study, showing their abbreviated name, sex, full name, culture, name of the original data set / publication, age, country of origin, and coordinates (latitude and longitude).

**Data S2. Genotypes retrieved from each individual**. Genotypes are shown for each of the SNPs from DisGeNET, and each individual analyzed. This table also shows the quadrant, time period, and DisGeNET code associated with each SNP.

**Data S3. List of risk alleles**. Risk alleles are shown for each of the SNPs from DisGeNET, as retrieved from SNPedia and SelfDecode.

**Data S4. Alleles frequencies**. Each subtable pertains to a particular quadrant (Q1 to Q4), and shows a list of DisGeNET SNPs, with their associated frequency in each time period, along with DisGeNET disease identifiers, full names, DisGeNET scores, number of records in PubMed, source, and CADD score.

